# Health service utilisation lags behind maternal diseases or illnesses during pregnancy in rural south Ethiopia: A prospective cohort study

**DOI:** 10.1101/593525

**Authors:** Moges Tadesse, Eskindir Loha, Kjell Arne Johansson, Bernt Lindtjørn

**Affiliations:** Dilla University, Ethiopia; Hawassa University, Ethiopia; Department of Global Public Health and Primary Care, University of Bergen, Norway; Centre for international health, University of Bergen, Norway

**Keywords:** Maternal, Illnesses, Health-service-utilisation, Cohort, Rural, Ethiopia

## Abstract

Maternal survival has improved substantially in the last decades, but evidence on maternal morbidity and health service utilisation for various maternal diseases are scarce in low resource settings. We aimed to measure health service utilisation for maternal illnesses during pregnancy. A cohort study of 794 pregnant women in rural southern Ethiopia was carried-out from May 2017 to July 2018. Disease or illness identification criteria were: symptoms, signs, physical examination, and screening of anaemia. Follow-up was done every two weeks. Data on health service utilisation was obtained from women and confirmed by visiting the health facility. Multilevel, multiple responses, repeated measures, and generalized linear mixed model analysis were used. The cumulative incidence of women experiencing illness episodes was 91%, and there were 1.7 episodes of diseases or illnesses per woman. About 22% of pregnant women were anaemic and 8% hypertensive. Fourteen pregnant women experienced abortions, 6 had vaginal bleeding, 48% pain in the pelvic area, 4% oedema, and 72% tiredness. However, health service utilisation was only 7%. About 94% of anaemic women did not get iron-folic-acid tablet supplementation. Only two mothers with blurred vision and severe headache were referred for further treatment. The main reasons for not using the health services were: the perception that symptoms would heal by themselves (47%), illness to be minor (42%), financial constraints (10%), and lack of trust in health institutions (1%). Risk factors were being older women, poor, having a history of abortion, living far away from the health institution, travelled longer time to reach a health institution, and monthly household expenditure >=30 USD. In Conclusion, there was a high incidence of diseases or illnesses; however health service utilisation was low. Poor understanding of severe and non-severe symptoms was an important reason for low health service utilisation. Therefore, community-based maternal diseases or illness survey could help for early detection. Ministry of Health should promote health education that encourages women to seek appropriate and timely care.

## Introduction

Maternal survival has improved substantially in the last decades, but evidence on maternal morbidity and use of health services for various maternal diseases or illnesses are scarce in low resource settings. For each woman who dies of maternal causes, at least 20 women suffer from maternal diseases or illnesses [1]. Thus, it is important to get more precise estimates of maternal disease burden in low-income settings [2]. Maternal diseases or illnesses lack common measurement indicators, standardized and agreed tools and methods of assessment at community or primary health care level [3]. Most of the earlier researches were institutional-based studies, and there are few population-based cohort studies that integrate diagnosis and self-report [4].

Though ensuring healthy lives and promoting the well-being of women is one of the agenda in the Sustainable Development Goal Number 3 [5], the disease or illness burden in developing countries among pregnant and women who gave birth remained high [6]. Globally, 10 to 20 million women suffer from pregnancy and childbirth-related diseases or illnesses annually [7]. In developing countries, 25% of pregnant women reported at least one episode of disease or illness during pregnancy [8]. About 14% of pregnant women in Ethiopia reported in 2013 at least one disease or illness during pregnancy [9].

Appropriate and timely care seeking and health service utilisation are essential for healthy living. During pregnancy, the use of health services is dependent on the women’s socioeconomic and demographic environment [10], like the mother’s age, educational status, occupation, access, and travel time to health facility [11].

Diseases during pregnancy are associated with both increased maternal deaths and stillbirths [12, 13]. Of the severe diseases in 2017, 56 million pregnant women experienced spontaneous abortion globally, and 49 millions of these were in low-income countries [14]. In Ethiopia, in 2017, the rate of spontaneous abortion was 28 per 1000 women [14]. In 2015, the global number of stillbirths was estimated to be 2.6 million [15] and Ethiopia accounted for 4% of these (97,000 stillbirths) [15]. Around 42% of pregnant women in developing countries were estimated to suffer from anaemia in 2017 [16], and 25% of Ethiopian pregnant women had anaemia [17]. Although iron-folic-acid supplementation is recommended for prevention or treatment of anaemia to all pregnant and women who gave birth [18], there is limited data about the uptake of iron-folic-acid supplementation during the postpartum period from Ethiopia [17]. Hypertension is also common during pregnancy, and 10 % of pregnant women globally [19], and 6% in Ethiopia had hypertension [20]. Hypertension is a well-known risk factor for maternal death, preeclampsia/ eclampsia, and cardiac and renal complications during pregnancy and the postpartum period [21].

As the causes and burden of maternal diseases or illnesses and use of health services in Ethiopia were not been well described in the literature, we conducted a cohort study on women attending antenatal care in a rural area of south Ethiopia with the aim to measure the incidence of diseases or illnesses during pregnancy, and simultaneously assess health service utilisation. Thus, our study presents new and important evidence for policies aiming at improving the health of mothers.

## Methods and materials

### Study design and period

We did a prospective cohort study on pregnant women attending antenatal care (ANC) in three local communities (kebeles) in south Ethiopia. The study lasted 15 months from May 2017 to July 2018.

### Study area and setting

This study was conducted in three randomly selected rural kebeles of Wonago district, in southern Ethiopia. Wonago district (zone) is found 420 km far away from the capital, Addis Ababa. It has 14 rural and 4 urban kebeles. A kebele is part of a district (wereda) which is the lowest administrative unit in Ethiopia and is composed of about 1000-1500 households (average 5000-7500 people) [22]. In 2017, the total population of the district was estimated to be 145,000 people [23]. The major ethnic group was Gedeo people, and the population density was 980 persons per km^2^. Agriculture is the dominant means of livelihood. The district has 6 health centres, 20 health posts, and 2 private clinics.

The three selected kebeles were Hase-Haro, Mekonisa, and Tumata-Chiricha in the Wonago district. The selected kebeles had 4 health posts, and 2 health centres and the total population was 28,822 people (9,420 in Hase-Haro, 13,304 in Mekonisa, and 6,098 in Tumata-Chiricha) [23]. The three kebeles were similar in socio-demographic and economic structure with most rural areas in Ethiopia. In 2013, more than 80% of the population lived in rural Ethiopia, and 26% of the rural residents lived on less than $1 per day, and 77% of rural mothers needed to travel more than 20 km to get to a hospital [24].

### Study participants

Pregnant women attending ANC formed the study population. After their second ANC visit, we followed each woman by visiting their homes every second-week until the mother gave birth.

### Inclusion and exclusion criteria

Pregnant women who were resident of the study area and had at least two ANC visits and were willing to participate in the study were included in the study. Those pregnant women who did not have any ANC visit, who did not reside in the study area and women who did not avail for examination at a scheduled visit were excluded after three attempts.

### Sample size

The sample size was determined by Openepi software Version 3.03 (www.openepi.com). To get the maximum sample size, we used different socio-demographic factors as exposure variables and the incidence of different types of diseases or illnesses during pregnancy as outcome variables. The assumptions were 95% level of confidence, 80% power, 1:1 ratio of unexposed to exposed in a sample, 41% of unexposed with an illness, and 1.5 relative risk. We estimated the sample size to be 898 pregnant women after adding 10% for non-response.

### Sampling procedure

First, we randomly selected 3 of 19 kebeles. Then, we recruited pregnant women attending the ANC in these kebeles until the required sample size was reached. We used house-to-house visits to follow-up the women until the women gave birth. There were 86 women with incomplete data, one withdrew from the study, and two women were excluded from the analysis as the women were confirmed as having cyst or mass rather than being pregnant which was decided in the nearest hospital (Fig. 1).

**Fig 1.**
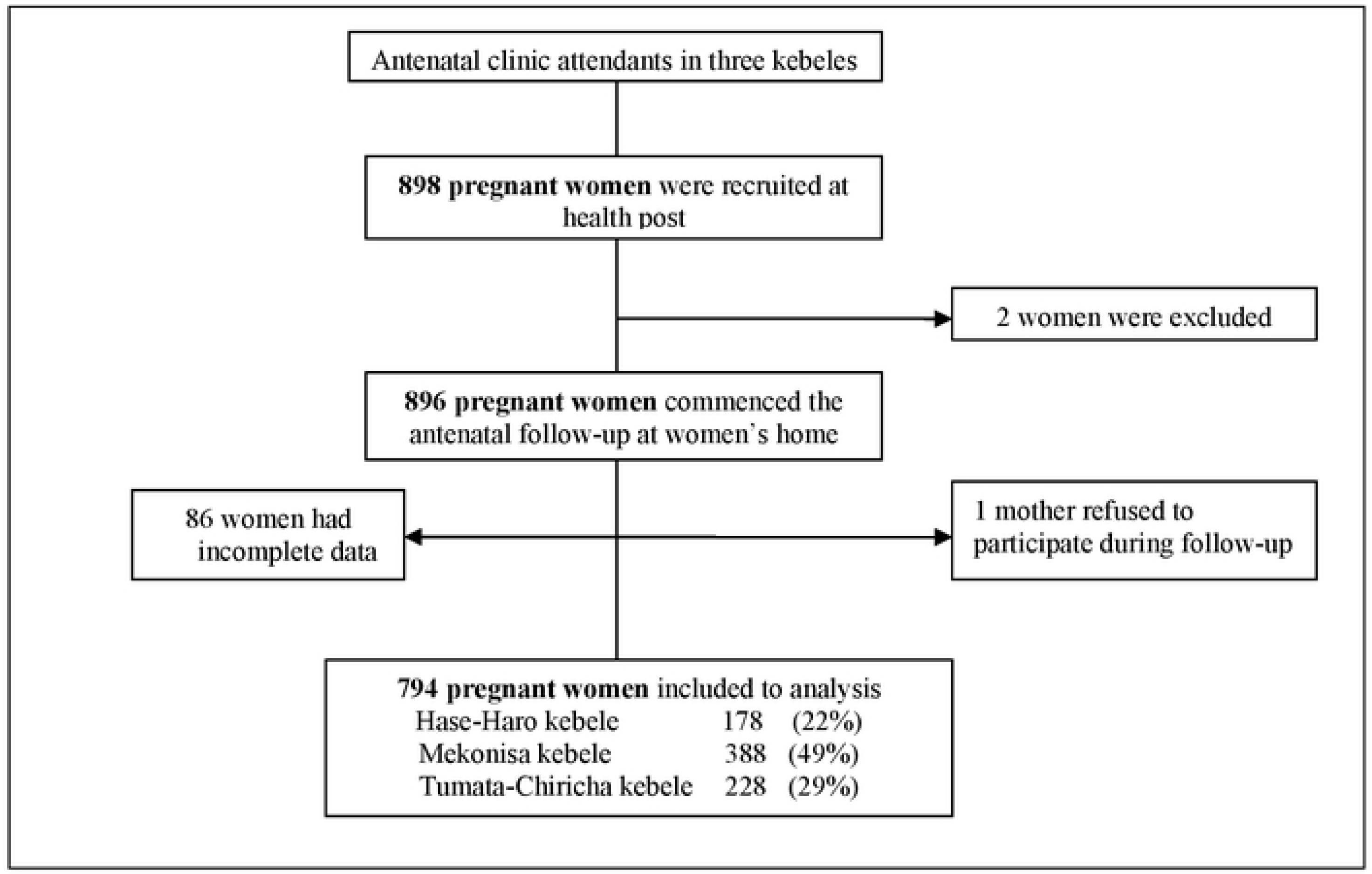
Flowchart of recruitment of pregnant women in rural southern Ethiopia, 2017/18

### Case definitions

The concepts of disease, illness, and sickness have often been used to indicate medical, personal, and social aspects of human ailments [25]. A woman with the disease was identified using disease or illness category and/or by the recording of an associated disability S1 Table. Disease or illness identification criteria were based on general symptoms, signs, physical examination, and screening of anaemia. Symptoms of maternal diseases or illnesses were used as they would correlate the associated disability, and maybe the primary reason for women to seek care. Signs were based on findings on general physical examination, and screening of anaemia was done for all women [2]. The obstetric diseases or illnesses were grouped as pregnancy termination, hypertension, haemorrhage, infection, and other diseases or illnesses. Medical diseases or illnesses were classified as gastrointestinal, psychiatric, and other diseases or illnesses [26].

As symptoms of diseases or illnesses during pregnancy are characterised as non-severe or physiological, mild, potentially life-threatening, and maternal near miss or deaths, we grouped the symptoms into categories based on their use of health services and on knowledge of which symptoms need health care such as bleeding and hypertension. Our assumption was that pregnant women would seek health care primarily because of medical problems. We then assessed how they used the health services for each of these groups.

### Operational definitions

Maternal diseases or illnesses: Any condition that complicates pregnancy and childbirth and which has a negative impact on a woman’s health and activity [2].

Disease: a medical or health problem that consists of physiological malfunction.

Illness: a personal or subjective undesirable state of health.

Sickness: the social aspects of human ailment [25].

Disability: a term for impairments, activity limitations and participation restrictions.

Stillbirth: a baby born with no signs of life at or after 28 weeks’ gestation [27].

Gravidity: all number of pregnancies in a lifetime (includes complete or incomplete).

Parity: number of children previously borne by a woman (excludes or abortions, but it includes stillbirths).

### Exposure variables

The exposure variables were socioeconomic and demographic factors which could affect the outcomes (incidence of maternal diseases or illnesses, and health service utilisation). Individual-level socioeconomic factors were: women’s age, marital status, women’s educational status, women’s occupation (housewife=0, others=1), household wealth, and total monthly household expenditure. Individual-level demographic factors were: women’s age at first marriage, women’s age at first birth, gravidity, parity, birth interval, history of abortion, and history of stillbirth. The community-level factors included their place of residence (Mekonisa=1, Hase-Haro=2, Tumata-Chiricha=3), type of road to the nearest health facility (asphalt=0, others=1), walking distance to the nearest health post or to the health centre or to hospital.

### Outcome variables

We used two sources of data to assess health service utilisation and the reasons for not seeking care. For each reported disease or illness episode, the women were asked whether they sought care or not, and then confirmed by visiting the health institutions. We recorded the symptoms of obstetric diseases or illnesses: hypertensive disorders, obstetric haemorrhage, pregnancy-related infection, and other diseases or illnesses. We also recorded symptoms of medical diseases or illnesses: gastrointestinal, psychiatric, and other symptoms.

The assessment for pregnancy-related disease or illness measurements were blood pressure and pulse rate (by Riester ri-champion^®^N digital apparatus, www.riester.de), and haemoglobin (HgB) (by HemoCue analyzer ®Hb 301 System, (www.hemocue.com). The validity and reliability of the measurements were checked regularly before used in the study.

Hgb values for pregnant women (3^rd^ Trimester, 27 weeks or more) was categorized into no anaemia (>=11 g/dl), mild anaemia (10–10.9 g/dl), moderate anaemia (7–9.9 g/dl), and severe anaemia (4–6.9 g/dl), very severe anaemia (<=3.9 g/dl) [28].

### Hypertension during pregnancy

Systolic blood pressure (SBP) was classified as normotensive (<=119.9 mmHg), pre-hypertensive (120-139.9 mmHg), and hypertensive (>=140 mmHg). Diastolic blood pressure (DBP) was defined as normotensive (<=79.9 mmHg), pre-hypertensive (80-89.9 mmHg), and hypertensive (>=90 mmHg) [29]. Hypertension was classified as either a systolic blood pressure of 140 mmHg or greater, or a diastolic blood pressure of 90 mmHg or greater, or both [30].

### Data collection

The data were collected prospectively using a pretested, structured, and an interviewer-administered questionnaire. The questionnaires were adopted from tools and indicators for maternal and newborn health [31], and from WHO maternal morbidity measurement working group [2]. The questionnaire was initially prepared in English and translated into local language, Gedeo language and Amharic, and back-translated into English. A pre-test was conducted among 82 mothers in a neighbouring kebele. The data collectors were trained women, residents of the selected kebeles who could speak the local language (Gedeo language) and Amharic and had completed grade 10. The field nurses and the supervisors were experienced in data collection and supervision. To ensure the data quality, double data entry and validation of data were employed using EpiData version 3.1 and analysed using SPSS version 25 software (SPSS Inc. Chicago, IL).

### Quantitative variables

The data were assessed using frequency distribution to explore if the variables were normally distributed. We used cross tabulation for categorical variables to see the distribution of each exposure variables on the outcome variables. Screening for missing values, outlier values, and data entry errors was done using frequency distribution and observation of the data. Errors were validated against the raw data and corrections were made before the analysis.

### Data analysis

Principal components analysis (PCA) was used to construct a wealth index of households based on 35 household assets and facilities (such as: type of houses’ roof, wall and floor, drinking water source, time, and safety, type and share of toilet facility, electricity/lighting, possessing radio, TV, mobile, telephone, watch, table, chair, bed, refrigerator, electric stove, and lamp, livestock, ox/milk cow, horse/donkey/mule, goat, sheep, chicken, beehives, place of cooking, having separate cooking room, plough, land ownership and size, saving, employment, and number of sleeping room). Before wealth index calculation was done, sample size adequacy was checked by Kaiser-Mayer-Olkin (KMO) value that should be >=0.6, Eigenvalue should be larger (>1) for factor 1 in order to rank the household by wealth status, and Bartlett’s test of sphericity should be significant, p-value <0.05 [32]. In this paper, the KMO value was 0.778, Eigenvalue was 3.97, and Bartlett’s test of sphericity p-value was 0.000. From the 35 questions used, 13 questions fulfilled this criterion. From 13 questions, 3 questions extracted with Eigenvalue above 1. By taking factor one, a ranking of cases was done. This wealth index was divided into two indices, and each household assigned to one of these indices (poor and rich).

Multilevel, multiple responses, repeated measures, and generalized linear mixed (GLMM) model analysis were used on the binary outcome variables to assess the individual and group processes associated with disease or illnesses and assessed the relationships between individuals (women) and their community (kebele). Four models were fitted using the GLMM in SPSS version 25. The model I (intercept-only model) was fitted without exposure variables to estimate the intraclass correlation coefficient (ICC) and to test the random variability of the intercept. Model II was examined for the effects of individual-level factors, Model III for community-level factors, and Model IV for both individual and community level factors sequentially [33].

ICC was calculated to evaluate whether the variation in disease or illnesses were primarily within or between individuals or communities. The Proportional Change in Variance (PCV) was obtained for each model against the null model to show the power of the factors in the model on disease or illnesses. It was calculated as PCV= (Ve-Vmi)/Ve; where Ve is variance in the null model and Vmi is variance in successive models. The study considered multilevel factors which might modify the effect of each other on disease or illness. The interaction effect of 20 exposure variables was checked and no significant interaction effects were seen. The Hosmer and Lemeshow recommendations were used in the selection of variables for a final model with a P-value 0.20 [34]. Statistical significance was considered at P-value <0.05. Multicollinearity and collinearity were checked among explanatory variables using mean-variance inflation factors (VIF).

### Ethical consideration

This study was approved by the Institutional Ethical Review Board at Hawassa University, College of Medicine and Health Sciences (IRB/100/08), and from the Regional Committees for Medical and Health Research Ethics (REC) of western Norway (2016/1626/REK vest). Written permission letters were obtained from the Gedeo Zone Health Department and the Wonago Wereda (district) health office. Written informed consent was obtained from each mother after she had received an explanation of the purpose of the study. The privacy, anonymity, and confidentiality of study participants were maintained. If a woman was found to have a disease or an illness, the nurses and data collectors linked the patient with the health extension workers in the kebele.

## Results

### Background characteristics

We obtained a response from 794 pregnant women (88.4%). The characteristics of the antenatal care population are listed in Table 1. From 794 pregnant women, 99.1% (787) were married, 63.7% (506) were illiterate, and 61.2% (486) was poor. The mean age was 25.3 years (range: 16, 45). The median walking distance to the nearest health post was 30 minutes (Inter Quartile Range, IQR: 20, 50), to the health centre it was 40 minutes (IQR: 25, 60), and to the hospital, it was 60 minutes (IQR: 40,180). The median monthly household total expenditure was 42.4 USD (IQR: 30.3, 60.1), of which 87% was spent on food. The mean age of pregnant women at their first marriage was 18.1 years (range: 14, 28), and the mean age at first birth was 19.4 years (range: 16, 28). About 23.3% (185) of mothers reported the current pregnancy was their first pregnancy, and 54.3% (431) had 2 or more children (Table 1). The proportion of observed to expected antenatal care visits was 71% from the three kebeles and varied from 49% to 96%.

**Table 1:** Characteristics of the antenatal care study population in Wonago district, rural south Ethiopia, May 2017 to July 2018 (=794)

| Socioeconomic characteristics |  | N | % |
| --- | --- | --- | --- |
| Kebele/residence | Mekonisa | 388 | 48.9 |
|  | Hase-Haro | 178 | 22.4 |
|  | Tumata-Chiricha | 228 | 28.7 |
| Age of mother in years | <20 | 177 | 22.3 |
|  | >=20 | 617 | 77.7 |
| Marital status | Married | 787 | 99.1 |
|  | Others | 7 | 0.9 |
| Maternal educational status | Not educated | 506 | 63.7 |
|  | Others | 288 | 36.3 |
| Maternal occupation | Housewife | 734 | 92.4 |
|  | Others | 60 | 7.6 |
| Wealth Index | Poor | 486 | 61.2 |
|  | Rich | 308 | 38.8 |
| Type of road to the nearest health facility | Asphalt | 208 | 26.2 |
|  | Others | 586 | 73.8 |
| Walking distance to the nearest health post | <=14 minutes | 114 | 14.4 |
|  | >=15minutes | 680 | 85.6 |
| Walking distance to the nearest health centre | <=29 minutes | 265 | 33.4 |
|  | >=30 minutes | 529 | 66.6 |
| Walking distance to the nearest hospital | <=59 minutes | 335 | 42.2 |
|  | >=60 minutes | 459 | 57.8 |
| Household total expenditure | <=30 USD/month | 18 | 2.3 |
|  | >=30.1 USD/month | 776 | 97.7 |
| Demographic variables |  |  |  |
| Age of mother at first marriage in years | <20 | 779 | 98.1 |
|  | >=20 | 15 | 1.9 |
| Age of mother at first birth in years | <20 | 697 | 87.8 |
|  | >=20 | 97 | 12.2 |
| Gravidity (No of pregnancy) | <=2 | 336 | 42.3 |
|  | >=3 | 458 | 57.7 |
| Parity (No of birth) | 1 | 334 | 42.1 |
|  | >=2 | 460 | 57.9 |
| Birth interval in years | <2 | 387 | 48.7 |
|  | >=2 | 407 | 51.3 |
| History of abortion | No | 746 | 94.0 |
|  | Yes | 48 | 6.0 |
| History of stillbirth | No | 756 | 95.2 |
|  | Yes | 38 | 4.8 |

1USD=27.64 ETB on August 31, 2018

### The incidence of diseases or illnesses and health service utilisation during pregnancy

The total disease or illness incidence was similar at all follow-up visits during pregnancy (46-53%), except during visit 1 where the incidence was highest (71.8%) (Table 2).

**Table 2:**
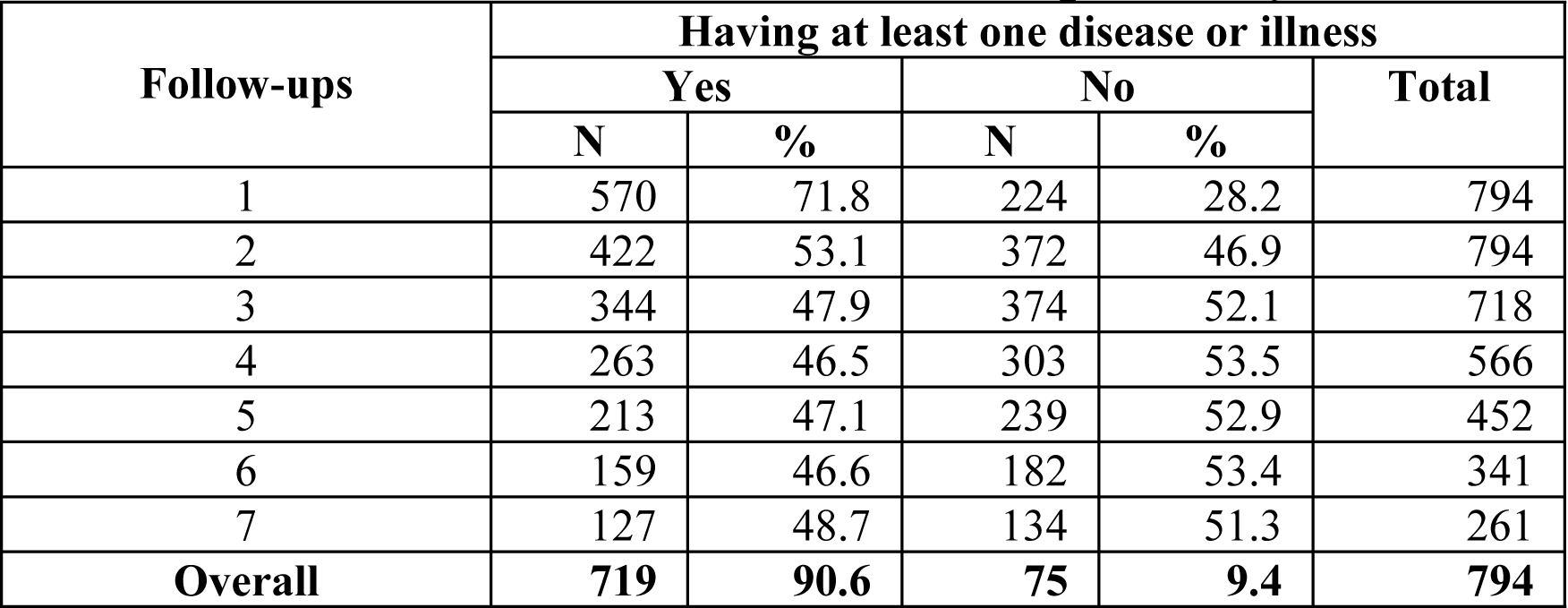
Distribution of pregnant women during follow-up and diseases or illnesses starting two-weeks preceding the first interview in rural southern Ethiopia, May 2017 to July 2018 (n=794)

| Follow-ups | Having at least one disease or illness |  |  |  | Total |
| --- | --- | --- | --- | --- | --- |
|  | Yes |  | No |  |  |
|  | N | % | N | % |  |
| 1 | 570 | 71.8 | 224 | 28.2 | 794 |
| 2 | 422 | 53.1 | 372 | 46.9 | 794 |
| 3 | 344 | 47.9 | 374 | 52.1 | 718 |
| 4 | 263 | 46.5 | 303 | 53.5 | 566 |
| 5 | 213 | 47.1 | 239 | 52.9 | 452 |
| 6 | 159 | 46.6 | 182 | 53.4 | 341 |
| 7 | 127 | 48.7 | 134 | 51.3 | 261 |
| Overall | 719 | 90.6 | 75 | 9.4 | 794 |

### Birth outcomes

About 64% (509) of women delivered at home, and of these, 96.3% (490) was attended by family members. There were 781 singleton and 13 multiple births. Vaginal deliveries accounted for 98.6% (783). Fourteen women experienced abortions (17.6 per 1000 pregnant women), and 26 deliveries resulted in a stillbirth (stillbirth rate 33.2 per 1000 births), and we observed one maternal death. None of the women who experienced abortion used health services.

The cumulative incidence of at least one type of disease or illness across all follow-up visits among 794 pregnant women was 90.6 (95% CI: 88.4, 92.4) per 100 pregnant-woman. There were 1.7 episodes of diseases or illnesses per woman, however, the overall rate of health service utilisation during disease or illness was only 6.7% (95%CI: 6, 7.4). The main reasons were symptoms that the women thought would heal by themselves in 46.4% (660 episodes), or that perceived illness was not important to seek health care that occurred among 42.2% (600), 10.3% (147 episodes) did not seek health care due to lack of money, and 1.1% (15) did not visit a health institution because they lacked confidence in the health institutions (Table 3).

**Table 3:**
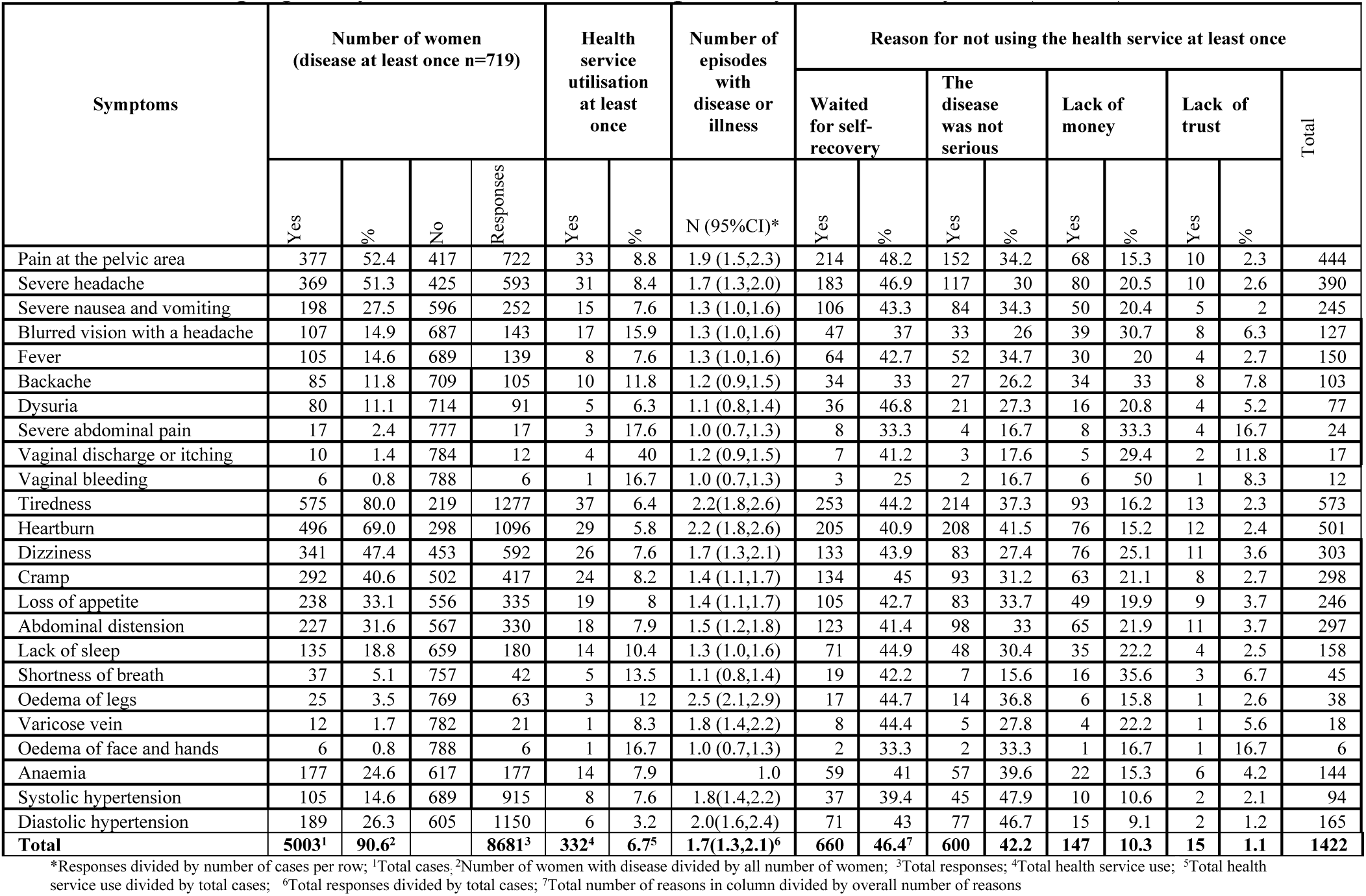
Symptoms of diseases or illnesses and reasons for not using the service during pregnancy in rural southern Ethiopia, May 2017 to July 201 (n=794)

Among those 719 pregnant women with the diseases or illnesses, the most common complaints were tiredness; 80% (95%CI: 77, 83). This was followed by heartburn, 69% (95%CI: 66, 72); pain at pelvic area, 52.4% (95%CI: 49, 56); and severe headache, 51.3% (95%CI: 48, 58). Others diseases were blurred vision with a headache, 14.9% (95%CI: 12, 18); fever, 14.6% (95%CI: 12, 17); 4.3% (95%CI: 3, 6) oedema; 0.8% (95%CI: 0.3, 1.7) vaginal bleeding; and foul smelling vaginal discharge, 1.4% (95%CI: 0.7, 2.5) (Table 3).

Among diseases that we regarded as severe and needed to be examined by health workers were six women with vaginal bleeding, of whom only one used the health service. Although 22% of pregnant women were anaemic, the uptake of Iron-folic-acid tablet supplementation was low (6%), and the reasons given were that 22 (15.3%) did not have money to visit a health institution, and 6 (4.2%) had no trust in the health institution. Their incidence of diastolic blood pressure was 26.3%; however, the rate of use of health service was only 3.2%. Among 31 (4.3%) women who had oedema, only 13% of them used the health service.

Among illnesses that we regarded as non-severe, there was low health service utilisation for tiredness, backache, and dizziness as many women with symptoms perceived their illness to be minor, and they thought they were not important, and some would heal by themselves (Table 3).

Only two of the mothers who had a severe headache and excessive tiredness or visual disturbance with a severe headache were referred for further treatment.

### Anaemia

Of 794 pregnant women, 22.3% (95%CI: 20, 25) were anaemic, of which, 13.9% (110) was mild, 7.9% (63) moderate, and 0.5% (4) was severe. However, 93.5% (95%CI: 92, 95) (742/794) of pregnant women did not get Iron-folic-acid tablet supplementation. But, the rate of use of health services during anaemia was only 7.9% (Table 3).

### Hypertension during pregnancy

Of 794 pregnant women, 7.6% (60) (95%CI: 6, 10) experienced hypertension. About 13.3% (105) of pregnant women had systolic, and 23.8% (189) had diastolic hypertension.

### Multilevel, mixed effect, repeated measures, and logistic regression analysis of maternal diseases or illnesses and health service utilisation

During disease or illness analysis, there was a model improvement and AIC increment from a model I to model IV. In the four models, the variability was significant between kebeles. The community-level variance (ICC) was 25.2% (95%CI: 19.4, 30.8 (Table 4). Risk factors for disease or illnesses include increasing age, poverty, history of abortion, time travel to a health facility, and if the pregnant mother lived far away from the health institution. The risk of disease or illness increases by 3.5% when the women’s age increases by one year and by 45% for women who were daily labourer and farmer. The pregnant women from households with monthly household expenditure >=30.1 USD were 61% less likely to have a disease. The pregnant women who had a history of abortion in their lifetime were 82% more likely to have an illness. In addition, the place of residence was a possible predictor of diseases or illnesses. Pregnant women living in Hase-Haro kebele were 52% less likely to have a disease or illness (Table 4).

**Table 4:**
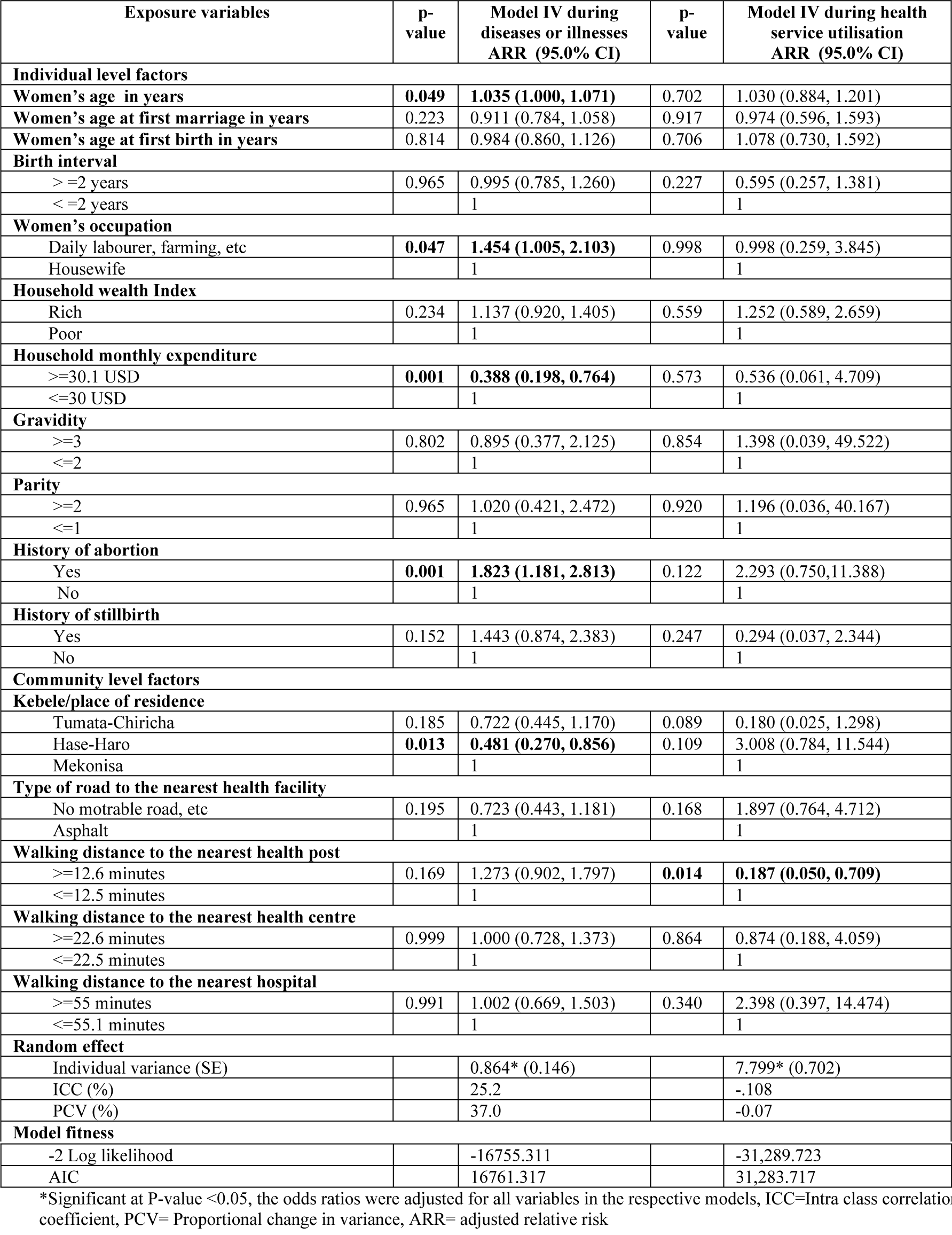
Multilevel mixed effect repeated measures logistic regression analysis of individual level and community level factors for diseases or illnesses and health service utilisation during pregnancy in rural southern Ethiopia, May 2017 to July 2018 (n=794)

However, during health service utilisation analysis, there was no model improvement from the model I to model IV, but AIC was increasing. The pregnant women who walked more than 15 minutes to reach the nearest health post were 81.3% less likely to use of health services when compared to the women who walked less than 15 minutes (Table 4).

## Discussion

In rural south Ethiopia, there was a high incidence of illness and disease among pregnant women and low health service utilisation. Over 90% of pregnant women experienced at least one symptom or disease during the period of observation. About 7.6% of pregnant women were hypertensive, and 22% were anaemic, and few of them received appropriate care. Risk factors for diseases or illnesses include increasing age, poverty, history of abortion, and if the pregnant mother lived far away from the health institution. Although many of these illness episodes were regarded and minor by the women, however, for potentially severe conditions such as hypertension, anaemia, and vaginal bleedings the rate of health service utilisation was low. The potential reasons for these results are discussed below:

In our study, the rate of diseases or illnesses was higher than previously conducted studies in Ethiopia [9], India [35], and in Pakistan [36]. This discrepancy could be explained by the fact that our study was a cohort study with multiple visits to pregnant women homes, and the other studies were cross-sectional. However, the incidence of diseases or illnesses was similar to 90.3% from population-based studies in Sri Lanka [37]. The Sri Lankan study showed that maternal illnesses considered as minor were judged not to be minor for women. Though, as shown by our study, the overall rate of use of health services during diseases or illnesses was low. The reasons for not seeking care during diseases or illnesses included a lack of money, or it could be due to longer time to travel to a health facility, or it could be that women who live in hard to reach areas with limited access to health facilities experienced an increased risk of diseases or illnesses than those who lived nearby to health facilities.

A ‘severe’ form of maternal diseases during pregnancy could include bleeding, anaemia, hypertension, and fever [38]. Women with these symptoms did not get the necessary care at the health posts and were not referred for treatment. Our study highlights that only 37 out of 582 women with bleeding, anaemia, hypertension, and fever used the health care service. However, a study from Egypt indicted that all mother with such diseases used the health services [39]. This difference could be due to the socioeconomic status of the women, some lacked trust in the health institutions, and some were not able to seek health care because of financial constraints(Table 3), and if the pregnant mother lived far away from the health institution. These findings emphasize that the need to identify and address barriers to health service utilisation, assess the quality of care, re-training of the health extension workers on disease or illness identification and take necessary public health measures.

The burden of anaemia in our study was similar to other studies in Ethiopia [40, 41], and it was most likely caused by iron deficiency [42]. However, our findings show that that the uptake of Iron-folic-acid tablet supplementation and health service utilisation for anaemia was low. These groups of pregnant women with anaemia were more likely to have an iron deficiency. This low health service utilisation during anaemia may put the women at risk as they are ‘severe’ as documented in our findings. The possible explanation could be poor women may prefer to remain at home may be due to a longer time to travel to health institution or if the pregnant mother lived far away from the health institution could avoid seeking care at all due to their inability to pay for health services and transportation and some due to lack of trust on the health institution.

In this study, the incidence of hypertension was high, however few of hypertensive women used the health service, as has also been reported from Bangladesh [43]. This finding indicated that the women who had hypertension could be at higher risk of complications of the mother and child, pre-eclampsia and eclampsia. In addition, only two of the mothers with the disease were referred for further treatment. This could be explained that women with these diseases were supposed to incur cost in order to get health care services (Table 3). Although health care services for pregnant women are free, however, if the women experience a disease or illness, they have to pay for health care and transportation. Therefore, these institutions should discuss how they could cover these costs.

A ‘non-severe’ form of maternal illnesses during pregnancy could include dizziness, abdominal cramp, headache, dysuria, shortness of breath, abdominal distension, lack of sleep, pain in the pelvic area and tiredness and it could affect the day-to-day life of pregnant women [37]. In this study, many pregnant women experienced pain in the pelvic area and tiredness and they did not use the health service. This finding was comparable with study in rural Bangladesh in which 87% of pregnant women had illnesses during their pregnancy, however, 73% of them did not seek any care [11]. This finding was also similar with Australian longitudinal study in which many women had non-severe illnesses, but 68% did not use health service [44], and in Sri Lanka, 90% did not use health service [37]. The use of health services is not primarily determined by a woman’s recognition of a problem. The possible explanation was that pregnant women with illnesses were reluctant to seek, and delayed in seeking health care as pregnant women with symptoms perceived their illness to be minor, and they thought they were not important, and some would heal by themselves. Further delay in seeking care may result from the perception that some women lack trusts in the health facilities and the underlying household poverty also plays a role in determining health care utilization as the pregnant women who could not afford for the use of health services.

In our study, we reported one of the study areas had higher risks than the other due to community factors. The variability due to community-level factors was significant between kebeles, as has been described from south-central Ethiopia [9]. This variability indicates that the presence of other factors which was not addressed in this study. The unexplained community-level variance remained significant and needs further study.

### Study design

the World Health Organization designed a template on how to estimate the incidence of diseases or illnesses during pregnancy based on a cross-sectional study design [2]. However, our results suggest that many diseases or illness episodes were not regarded as severe by the patients and further work on how to improve the questionnaires may be needed and one possibility is to ask about use of health services.

Our methods of registration of symptoms based on a recording of associated disability during pregnancy revealed that high incidence of maternal diseases or illnesses, however, there was low health service utilisation. Community-based disease or illness survey could be used for early detection, to improve the health of mothers, and take necessary public measures. In this regard, this study coincides with the World Health Organisation maternal morbidity matrix for assessing maternal diseases or illnesses [2]. The need to measure and count ‘severe’ or ‘non-severe’ disabilities reported by the women could help to address the health needs of women and could serve as an indicator of access to health care, quality of the health care system, and the possibility of a survey of maternal diseases or illnesses for an informed decision.

### Strengths of the study

To our knowledge, this study is the first of its kind in Ethiopia to study factors associated with diseases or illnesses during pregnancy and use of health services. It was an antenatal care based cohort study with multilevel, mixed effect, and repeated measures to identify the multilevel determinants of diseases or illnesses.

### Limitations of the study

In this study, only those pregnant women who attended antenatal care were recruited: This might be a potential selection bias in this study as a random selection of women from the communities was not employed. The proportion of observed to expected antenatal care visits in the three kebeles was in agreement with birth registry studies in Southwest Ethiopia where the coverage of maternal health services was about 75% [45]. Thus, our study may be representative of women attending antenatal care but may not be fully representative of women not attending such services. Women not attending antenatal care could have higher maternal incidence rates of severe diseases, and lower use of health services. As there was a displacement of residents in the study area and its surrounding [46], the number of pregnant in the analysis was 88.4% due to incomplete data. Despite these limitations, this study provides useful insights about the incidence of diseases or illnesses and use of health services and it would serve as a base for more detailed investigation.

### The policy implication of the findings

In conclusion, maternal disease or illnesses during pregnancy was high and health service utilisation was low. Poor understanding of what severe and non-severe symptom was an important reason for low health service utilisation. Therefore, community-based maternal disease or illness survey could help for early detection. Ministry of health should promote health education that encourages women to seek appropriate and timely care, which in turn could improve the use of health services. We recommend further an interventional study should be carried out to answer why health service utilisation was low and how it could be improved, and how diseases or illnesses could be prevented and how the health of mothers could be improved.

## Acknowledgements

We would like to acknowledge Hawassa University, College of Health Sciences and Medicine, School of Public Health, and the University of Bergen, Centre for International Health, for providing the opportunity to do this study. My deepest gratitude also goes to SENUPH/ NORHED project (*The* South Ethiopia Network of Universities in Public Health project/The Norwegian Programme for Capacity Development in Higher Education and Research for Development) for giving me the chance to join PhD programme. Finally, we would also like to thank the mothers for their volunteer participation.

## Supporting Information

***Table S1:*** Sign and/or symptoms of diseases or illnesses and case definition

